# Experimental assessment of K locus effects on the gray wolf response to virus

**DOI:** 10.1101/2020.12.29.424723

**Authors:** Rachel A. Johnston, James G. Rheinwald, Bridgett M. vonHoldt, Daniel R. Stahler, William Lowry, Jenny Tung, Robert K. Wayne

## Abstract

In North American gray wolves, black coat color is dominantly inherited via a three base pair coding deletion in the *canine beta defensin 3* (*CBD103*) gene. This three base pair deletion, called the *K^B^* allele, was introduced through hybridization with dogs and subsequently underwent a selective sweep that increased its frequency in wild wolves. Despite apparent positive selection, *K^BB^* wolves have significantly lower fitness than wolves with the *K^yB^* genotype, even though the two genotypes show no observable differences in black coat color. Thus, the *K^B^* allele is thought to have pleiotropic effects on as-yet unknown phenotypes. Given the role of skin-expressed *CBD103* in innate immunity, we hypothesized that the *K^B^* allele influences the gene regulatory response to viral infection. To test this hypothesis, we developed a panel of primary epidermal keratinocyte cell cultures from 24 wild North American gray wolves (both *K^yy^* and *K^yB^* genotypes) and generated immortalized *K^yy^* and *CBD103* knockout lines. We assessed the transcriptome-wide responses of wolf keratinocytes to polyinosinic:polycytidylic acid (polyI:C), which mimics infection by a double-stranded RNA virus, and to live canine distemper virus. Keratinocytes with the *K^yB^* genotype and with the *K^yy^* genotype had similar gene regulatory responses to viral infection, suggesting that this response does not explain pleiotropic effects of the *K^B^* allele on fitness. This study supports the feasibility of using cell culture methods to investigate the phenotypic effects of naturally occurring genetic variation in wild mammals.

## Introduction

In mammals, much of the variation in pigmentation traits, especially fur and hair color, is controlled by the agouti-melanocortin receptor 1 (MC1R) pathway (Nordlund *et al.*, 2008). North American gray wolves exemplify this pattern: coat color can be gray or black, where black coat color is controlled by a dominant three base pair deletion in the coding sequence of the gene *canine beta defensin 3* (*CBD103*) (Figure 1A) (Candille *et al.*, 2007). This coding deletion, known as the *K^B^* allele, increases the binding affinity of the CBD103 protein to MC1R. CBD103-MC1R binding in turn competitively excludes the Agouti protein from binding to MC1R, shifting the balance of pigment production to the dark pigment eumelanin; the same set of interactions explains, in part, coat color polymorphism in domestic dogs (Candille *et al.*, 2007).

**Figure 1.**
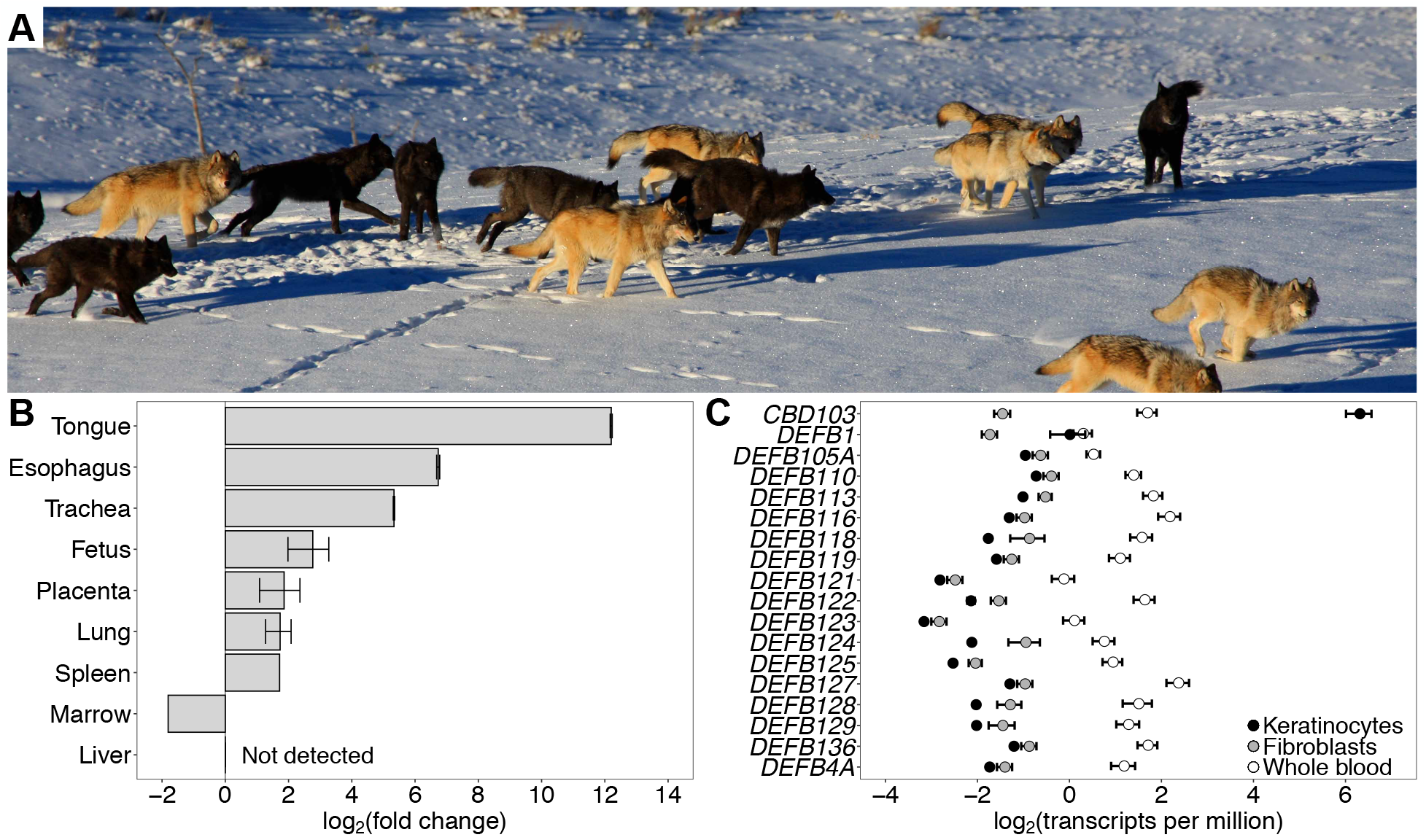
*CBD103* gene expression in North American gray wolves. **(A)** Coat color polymorphism in North American gray wolves in Yellowstone National Park, in which coat color can be gray or black (photo credit Dan Stahler/National Park Service photo). Black coat color is dominantly inherited, conferred by a three base pair coding deletion in *canine beta defensin 3* (*CBD103*). **(B)***CBD103* expression, relative to expression in dog testes, across tissues of a recently deceased pregnant female *K^yy^* wolf. Error bars represent standard errors across RT-qPCR replicates (2 – 3 replicates per tissue). A single tissue sample was collected for each tissue except fetus and placenta, which each represent two tissue samples (i.e., from two fetuses). **(C)** Absolute expression of annotated beta defensins in epidermal keratinocytes (N = 23), fibroblasts (N = 6), and whole blood (N = 25; (Charruau *et al.*, 2016)) from North American gray wolves. Only *CBD103* is highly expressed in keratinocytes.

In wolves, the *CBD103* polymorphism is notable not only for being one of the few well-understood genotype-phenotype relationships in wild mammals, but also because of its adaptive importance. The deletion allele was introduced to North American gray wolves from domestic dogs through hybridization circa 1,000-10,000 years ago, likely with dogs brought with Native Americans. After it was introduced, it subsequently underwent a selective sweep, either due to effects on coat color or pleiotropic effects on other phenotypes (Anderson *et al.*, 2009; Coulson *et al.*, 2011; Schweizer *et al.*, 2018). However, despite evidence of positive selection for the *K^B^* allele (Anderson *et al.*, 2009; Schweizer *et al.*, 2018), individuals that are homozygous for the *K^B^* allele (*K^BB^*) experience lower annual recruitment and survival rates and much lower mean lifetime reproductive success than *K^B^* allele heterozygotes (*K^yB^*, where the wild-type allele is conventionally abbreviated *K^y^* for “yellow”). Moreover, in the well-sampled Yellowstone National Park population, fewer *K^BB^* individuals have been detected than expected based on observed allele frequencies (Stahler, unpublished data). The reduced fitness of *K^BB^* individuals relative to *K^yB^* individuals, as well as extensive population genetic simulations, thus suggest that both *K^B^* and *K^y^* alleles are currently being maintained in wild wolves as a consequence of balancing selection (Coulson *et al.*, 2011; Schweizer *et al.*, 2018).

The phenotypic costs of *K^B^* homozygosity remain unclear. Although the coat color phenotype is the most readily apparent consequence of the *K^B^* allele, both *K^BB^* and *K^yB^* wolves exhibit the same black coat color (Coulson *et al.*, 2011). The low recruitment of *K^BB^* animals, and resultant balancing selection, may be explained in part by disassortative mating, in which wolf pairs that successfully produce offspring tend to include one parent with black, and one parent with gray, coat color (Hedrick *et al.*, 2016). However, the apparent survival and lifespan differences between *K^yB^* and *K^BB^* animals, and rarity of *K^BB^* individuals even from pairs of *K^yB^* parents, suggest that the *K^B^* allele also confers as-yet unknown pleiotropic effects in North American gray wolves that account for markedly lower fitness in *K^B^* homozygotes (Coulson *et al.*, 2011; Schweizer *et al.*, 2018). Indeed, pleiotropy of *K^B^* is indirectly supported by the reported effects of coat color on aggression (Cassidy *et al.*, 2017) and on pup survival (Stahler *et al.*, 2013).

The most obvious class of candidate phenotypes is immune defense. Defensin genes are named for their role in pathogen defense, immune signaling, and generation of antimicrobial peptides (García *et al.*, 2001; Harder *et al.*, 2001; Ganz, 2003; Wilson *et al.*, 2013). The human homolog of *CBD103*, *DEFB103*, is expressed in epithelial cells (especially tonsil, skin, trachea, and tongue) as a first line of defense against pathogens (García *et al.*, 2001; Harder *et al.*, 2001; Wilson *et al.*, 2013). For example, its protein product, HBD3, is known to inhibit herpes simplex virus infection by deterring binding and entry of the virus into cells (Hazrati *et al.*, 2006), and inhibits human immunodeficiency virus (HIV) replication via direct binding with virions (Quiñones-Mateu *et al.*, 2003). Similarly, in domestic dogs, *CBD103* is expressed in skin and tongue (particularly in keratinocytes), and the CBD103 protein shows potent antibacterial activity (Candille *et al.*, 2007; Leonard *et al.*, 2012). Based on antimicrobial assays against five gram-negative and gram-positive bacterial taxa, antibacterial activity of CBD103 shows no clear differentiation between the *K^y^* and *K^B^* forms (Leonard *et al.*, 2012). However, the impacts of *CBD103 K^B^*/*K^y^* genotype on other aspects of immunity, such as antiviral activity, are not yet known.

In the present study, we tested the hypothesis that the gray wolf *K^B^* allele has pleiotropic effects on immunity via influencing the gene regulatory response to viral infection. To do so, we first identified the tissue and cell types that show the highest expression of *CBD103* in North American gray wolves. We then established a panel of primary epidermal keratinocyte cultures from a genetically diverse set of wild gray wolves (14 *K^yy^* and 10 *K^yB^*; note that *K^BB^* homozygotes are rare due to their large fitness disadvantage). Based on these cultures, we challenged both primary and immortalized wolf keratinocytes with polyinosinic:polycytidylic acid (polyI:C), which stimulates Toll-like receptor 3 to mimic infection by a double-stranded RNA virus. We also challenged a subset of the wolf keratinocyte cultures with a live, ecologically relevant pathogen, canine distemper virus (CDV), which is thought to be an important pathogen in explaining temporal spikes in wild wolf mortality (Almberg *et al.*, 2009). If the *CBD103 K^B^* allele alters the response to CDV or to general viral infection, then such an effect could contribute to the complex, unexplained relationship between *CBD103* genotype and host fitness. Finally, we demonstrated the utility of establishing new cell line resources from wild mammals by immortalizing *K^yy^* wild-type keratinocyte cells and using CRISPR/Cas9 gene editing to create a targeted *K^y^* to *K^B^* mutation. For the primary, immortalized, and edited cell lines, we then evaluated whether *CBD103* genotype predicts the gene regulatory response to viral infection.

## Materials and Methods

### Cross-tissue evaluation of CBD103 expression

To identify the wolf tissue with the highest *CBD103* expression, we leveraged opportunistically sampled tissue biopsies from a recently deceased *K^yy^* female wolf (animal ID: 759F; tissues collected < 48 hours from time of death). A small piece of each tissue (< 1 cm^2^ each of bone marrow, esophagus, fetus, liver, lung, placenta, skin, spleen, tongue, and trachea) was transferred with sterile forceps into a PAXGene Blood RNA tube and shipped to the University of California-Los Angeles (UCLA) on dry ice.

Total RNA was extracted from tissues using the Trizol Plus RNA Purification Kit with DNAse treatment (RNAse-Free DNAse Set, Ambion) and column cleanup (PureLink RNA Mini Kit, Invitrogen). For quantitative RT-qPCR, complementary DNA (cDNA) was synthesized using SuperScript II Reverse Transcriptase (ThermoFisher Scientific) on 200 ng RNA of each sample following the manufacturer’s instructions. The cDNA products were then purified using the QIAquick PCR Purification Kit (Qiagen). RT-qPCR reactions were performed on 10 ng cDNA using LightCycler 480 SYBR Green Master Mix (Roche) with 10 ul reaction volumes, using previously published *CBD103* and *rps5* primers and cycling conditions (Leonard *et al.*, 2012). The following program was used on a LightCycler 480 Instrument: 95°C for 5 min; 45 cycles of 95°C (10 sec) and 58°C (45 sec); subsequent melting curve that ramps the temperature from 72°C to 95°C in increments of 0.1°C/sec. RT-qPCR values were standardized to the expression level of *rps5*.

### Collection of skin biopsies for primary cell culture

Our cross-tissue survey indicated that *CBD103* was highly expressed in epithelial tissues (see Results), which are most readily sampled in wolves via skin biopsies from the ear. To collect skin biopsies from the field in a manner that allowed us to establish primary cell cultures, we pre-prepared culture medium nutrients in single sample aliquots (“MEFcubes”), which were stored frozen until use. Each MEFcube was comprised of 1 ml bovine calf serum (BCS; Hyclone, Cytiva Life Sciences), 0.1 ml L-glutamine (ThermoFisher Scientific), 20 ul Primocin antibiotic (Invivogen), and 252 ul 1 M HEPES buffer (ThermoFisher Scientific), and stored frozen up to 1 year. We also pre-prepared and refrigerated aliquots of 9 ml Dulbecco’s Modified Eagle’s Medium (DMEM; ThermoFisher Scientific) in 15 ml conical tubes (up to 1 year refrigeration). 0 – 3 days prior to tissue collection, collection medium was prepared for each sample by thawing a MEFcube and transferring it to a 15 ml tube containing 9 ml DMEM, and refrigerating the freshly prepared medium until the day of tissue collection.

Study subjects were anaesthetized prior to collection of a 6 mm skin biopsy by the National Park Service following protocols approved by the National Park Service Institutional Animal Care and Use Committee (Permit IMR_YELL_Smith_wolves_2012) and the Idaho Department of Fish and Game following United States national guidelines for handling animals. Upon collection, each 6 mm skin biopsy was immediately transferred to a 15 ml tube of collection medium and kept in a chilled cooler (Tovar *et al.*, 2008). A subset of biopsies were collected into 1 ml aliquots of collection medium (in 1.7 ml Eppendorf tubes) during field work instead of directly into a 15 mL conical tube and transferred to 15 mL tubes containing 10 ml collection medium at the end of the field day. Samples were then transferred to a refrigerator until overnight shipment to UCLA on wet ice.

### Primary culture of wolf keratinocytes and fibroblasts

Working in a Biosafety Class II cabinet, each skin biopsy was transferred to a p100 Petri dish containing 50 ul collection medium. The biopsy was then minced into *ca*. 0.5 mm pieces using curved iris scissors and evenly dispersed across the dish. A small drop of medium (*ca*. 20 ul) was placed around each skin piece, and 2 – 3 ml medium was added around the bottom of the dish’s rim. Samples were then incubated at 37°C under 5% CO_2_ and atmospheric oxygen levels. The medium was changed every 2 – 3 days. We evaluated four culture medium formulations for their promotion of wolf keratinocyte cell growth at initial plating (Supplementary Materials and Methods; Supplementary Figures S1-S3). From those results we used the following optimized medium (Wu *et al.*, 1982) (hereafter referred to as “FAD medium”) to initiate primary keratinocyte cultures from the biopsies: 1:1 DMEM:F12 base media (ThermoFisher Scientific) + 5% BCS + 0.4 ug/ml hydrocortisone + 10 ng/ml epidermal growth factor (EGF) + 1% Penicillin-Streptomycin (ThermoFisher Scientific). After two days incubation, 10 uM ROCK inhibitor Y-27632 (hereafter referred to as RI) [Cayman Chemical] was added (Liu *et al.*, 2012). All images of cells were taken using an AxioCamMRm camera (Zeiss).

To separate fibroblasts from keratinocytes after initial plating (*ca*. 1 week post-plating), dishes were first washed once with 1x phosphate buffer solution (PBS; ThermoFisher Scientific). 1 ml 0.25% trypsin-EDTA (ThermoFisher Scientific) was then added to each well and quickly pipetted up and down 3 – 5 times, which detached fibroblasts but not keratinocytes. This initial cell suspension, containing fibroblasts, was transferred to a 15 ml tube containing 1 ml DMEM + 10% BCS. The dish was then quickly rinsed two times with PBS to remove any residual fibroblasts and then was incubated with 1 ml 0.25% trypsin-EDTA at 37°C for 5 – 10 minutes to detach keratinocytes for cell passage. After this initial separation, selective growth of each cell type was performed using cell type-specific media. Upon passage of cells, we compared 12 growth treatments for culturing primary wolf keratinocytes (see Supplementary Materials and Methods). We found that primary wolf keratinocytes grow best in FAD medium (formula above) + RI, with 3T3 feeder cells (Todaro and Green, 1963), which are known to promote growth of human epidermal keratinocytes (Rheinwald and Green, 1975) (see Supplementary Materials and Methods). We also cultured the separated fibroblasts, growing them in M199/M106 medium, a formulation optimal for human fibroblasts [(Dickson *et al.*, 2000): 1:1 (vol/vol) medium M199 and medium M106 (ThermoFisher Scientific) + 15% BCS + 10 ng/ml EGF + 0.4 ug/ml hydrocortisone + 1% Penicillin-Streptomycin], to permit quantification of *CBD103* expression levels in this cell type.

Cells were cryopreserved in DMEM/F12 + 10% BCS + 10% dimethyl sulfoxide (DMSO; Sigma) by transferring cryotubes to a room temperature Mr. Frosty Freezing Container (ThermoFisher Scientific) which was placed in a −80° C freezer overnight. Cryotubes were then transferred to a liquid nitrogen freezer for storage.

### Immortalization of wolf keratinocytes and generation of a K^yB^ cell line using CRISPR/Cas9

Wildtype *K^yy^* cells were immortalized and edited to generate a *K^yB^* line by Applied StemCell, Inc (Milpitas, CA, USA). Generation of a *K^BB^* line was also attempted, but was not successful. Keratinocytes from a single *CBD103* wildtype *K^yy^* individual (animal 15071) were first immortalized by infecting 1 × 10^5^ cells with freshly packaged lentivirus encoding simian virus 40 (SV40) T antigen following Applied StemCell’s standard protocol. The cells were selected with puromycin and passaged for 5 – 8 passages. SV40 transgene expression of puromycin-selected cells was determined by PCR amplification of a 112 base pair amplicon of SV40 using the following primers: SV40-F: 5’-GGGAGGTGTGGGAGGTTTTT-3, SV40-R: 5’-TCAAGGCTCATTTCAGGCCC-3’. The resultant wolf immortalized keratinocyte (WIK) cell line was subsequently cultured in DMEM containing 10% fetal bovine serum, 1x non-essential amino acids, 1x L-glutamine, 1x sodium pyruvate, and 1% Penicillin-Streptomycin. Immortalized cells were plated at 2 × 10^4^ cells per well of a 6-well plate. Medium was changed every day, and cells were passaged every 4 – 5 days or when confluent.

To generate the *K^B^* allele, which is the deletion of the three base pairs TCC/AGG at chr16:58965448-58965450 (*canfam3.1*), the following guide RNA (gRNA) sequence was selected based on proximity to the target three base pair deletion: CCTGCAGAGGTATTATTGCAGA. To prevent the gRNA/Cas9 complex from recognizing and cutting sequence after the single-stranded donor oligonucleotide (ssODN) was used as a repair template, a silent single nucleotide polymorphism (SNP), corresponding to the first base pair of the gRNA, was incorporated into the ssODN (C to T). Additionally, an intronic SNP (C to G) was introduced by the ssODN at chr16:58965507, as the dog genome was used to design the ssODN and is fixed for G at that genomic coordinate.

To screen clones for the three base pair deletion, the following primers were developed to amplify a 298 base pair region containing the three base pair *CBD103* deletion and two TspEI digestion sites: forward 5’-GTGAGGTGTACAATGAGGATTATAACTGAACTCC, reverse 5’-GGAAGAACAGCGGCCTATCTGC. TspEI digestion of the PCR product of *K^y^* yielded three fragments of lengths 112 base pairs, 103 base pairs, and 83 base pairs, whereas TspEI digestion of the PCR product of *K^B^* yielded two fragments, of lengths 112 base pairs and 183 base pairs (Supplementary Figure S4). Clones thought to carry the intended three base pair deletion were sequenced with Sanger sequencing. Sanger sequencing suggested that one clone (clone ID: P2H9) was heterozygous for the intended three base pair deletion.

To confirm K locus genotype and characterize the immortalized *K^yy^* cell line and the *K^yB^* edited cell line, we performed whole genome sequencing. We extracted DNA from each cell line using the DNeasy Blood and Tissue Kit (Qiagen) and prepared DNA-Seq libraries from 100 ng DNA using the NEBNext Ultra II FS DNA Library Prep Kit for Illumina (New England Biolabs; 12 minutes enzymatic shearing, 4 PCR cycles). 100 base pair, paired end sequencing was performed on a NovaSeq SP at the Duke Sequencing and Genomic Technologies Shared Resource core facility. Reads were trimmed with Trim Galore (Krueger, 2015) to remove adapter sequence and base pairs with Phred score < 20 from the ends of reads, keeping only reads ≥ 20 basepairs long after trimming. We mapped reads to the dog genome (*canfam3.1*) using *bwa* with default settings and removed reads with MAPQ less than 10 (Li and Durbin, 2009). We performed indel realignment and base recalibration using the Genome Analysis Toolkit (GATK) (McKenna *et al.*, 2010). Base recalibration was based on genotypes with GQ ≥ 4 from initial genotyping of the data with the UnifiedGenotyper function. Final genotyping was performed with HaplotypeCaller. We removed variants that did not pass the following thresholds: QUAL < 100, QD < 2, MQ < 35, FS > 30, HaplotypeScore > 13, MQRankSum < −12.5, ReadPosRankSum < −8. Variants were then filtered to only retain biallelic variants with a genotype quality of at least 20, with ≥30x coverage in both cell lines. From the filtered dataset, we defined potential off-target edits as variants that had been called in the *K^yB^* edited cell line that had no reads supporting that variant in the parent immortalized *K^yy^* cell line.

### PolyI:C and live CDV challenge experiments

Keratinocytes (N = 48 samples, two samples per individual/cell culture) were plated at 1.5 × 10^4^ cells per well of a 12-well plate with 3T3 feeder cells in FAD + 10 uM RI. When cells were at *ca.* 30% confluence, they were switched to a low-serum medium (FAD + 0.5% BCS) without RI and incubated for 24 hours. Cells were then treated with 1 ug/ml polyI:C (Sigma-Aldrich) or vehicle control (sterile, endotoxin-free water) for 24 hours. Wells were quickly washed once with 0.25% trypsin (by pipetting up and down 3 times) and twice with PBS to remove 3T3 cells before collecting keratinocytes into 1 ml Trizol (Invitrogen). Details of the samples used for polyI:C challenge across the 24 wolf keratinocyte cultures are provided in Supplementary Table S1.

Canine distemper virus (CDV) challenges were performed on a subset of six primary keratinocyte cultures (from three *K^yy^* and three *K^yB^* animals; N = 12 samples, two samples per individual/cell culture) and on the *K^yy^* and *K^yB^* immortalized cell lines using CDV 5804PEH-eGFP, a recombinant wild-type CDV that expresses green fluorescent protein (GFP) (Von Messling *et al.*, 2004). CDV was propagated in VerodogSLAMtag cells (Von Messling *et al.*, 2004), and TCID_50_/cell was calculated following the Spearman-Kärber method (Finney, 1952). Keratinocytes were plated in duplicate on 24-well plates at 8 × 10^4^ cells per well in FAD + 5% BCS + RI, without feeder cells. After a 24 hour incubation, the medium was changed to FAD + 0.5% BCS. After an additional 24 hours, cells were infected at an MOI of 20 TCID_50_/ml or 100 TCID_50_/ml. At varying time points (5 days post-infection for primary cells, and 0, 1, 2, 3, 4, and 5 days post-infection for the immortalized cell lines), cells were collected in 1 ml Trizol for RNA preservation. Total RNA was extracted from cells using the Trizol Plus RNA Purification Kit with DNAse treatment (RNAse-Free DNAse Set, Ambion) and column cleanup (PureLink RNA Mini Kit, Invitrogen). For the primary and immortalized keratinocytes, images of CDV-infected cells (which express GFP) were captured using an AxioCamMRm camera (Zeiss).

### RNA-Seq data generation and analysis

We used RNA-seq to quantify *CBD103* gene expression in primary keratinocytes and fibroblasts, and to profile the immune response to polyI:C and CDV in the primary keratinocyte cultures and the *K^yy^* and *K^yB^* immortalized cell lines. RNA-seq libraries were generated on high quality RNA extracts (RNA Integrity number [RIN] was measured for the 48 polyI:C and control samples; mean = 9.83 +/− 0.52 s.d, Supplementary Table S1) using half-reactions from the TruSeq RNA Sample Preparation Kit (Illumina), following the manufacturer’s instructions. RNA-seq libraries were indexed and pooled at 12 samples per lane and sequenced as 100 base pair single-end reads on the Illumina HiSeq 4000 at the Vincent J. Coates Genomics Sequencing Laboratory at University of California-Berkeley.

The resulting reads (mean 29.62 million per sample +/− 4.65 million s.d.; Supplementary Table 1) were trimmed with Trim Galore (Krueger, 2015) to remove adapter sequence and base pairs with Phred score < 20 from the ends of reads, keeping only reads ≥ 20 basepairs long after trimming. Reads were mapped to the dog genome (*canfam3.1*) using two-pass mapping with STAR (Dobin *et al.*, 2013). Only uniquely mapped reads were retained. We quantified gene expression using HT-Seq (Anders *et al.*, 2014) with the “union” mode using the Canis_familiaris.CanFam3.1.94.gtf file from Ensembl. We transformed the raw counts to transcripts per million (TPM) (Wagner *et al.*, 2012) and removed genes that had an average TPM < 2 in either condition within each RNA-Seq data set (i.e., null or polyI:C; null or CDV). For each data set (the polyI:C data set and the CDV data set), we also removed the 5% of genes with the lowest variance within each condition. We performed *voom* normalization (Law *et al.*, 2014) within each data set using the voomWithQualityWeights function, with the trimmed mean of M-values (TMM) method (Robinson and Oshlack, 2010) in DESeq (Anders *et al.*, 2014) to estimate normalization factors. For the data set of primary keratinocytes challenged with polyI:C, we controlled for the technical effects of RNA-Seq library yield, sample provider (Yellowstone National Park Service or Idaho Department of Fish and Game), and PC2 (which improved power to detect *cis*-eQTL) by regressing them out from the expression level data for each gene using *limma* (Smyth, 2005). Two samples (polyI:C treated 1005F, and null control 14387) did not cluster in principal component space according to condition (null versus polyI:C-treated) and were removed from downstream analyses.

We modeled variation in the expression levels of each gene using mixed effects models fit in *emmreml* (Akdemir and Okeke, 2015). For each gene in each data set, we first ran a model in which normalized, batch-corrected expression levels were modeled as a function of the fixed effect of condition (null or stimulated) and a random effect to account for samples from the same individual and genetic relatedness across individuals (see Supplementary Materials and Methods for estimation of the relatedness matrix *K*). We then ran two additional models: 1) gene expression as a function of the fixed effects of condition and *CBD103* genotype nested within condition, and a random effect to control for repeated samples and genetic relatedness; and 2) gene expression as a function of the fixed effects of condition, *CBD103* genotype, and the interaction of condition and *CBD103* genotype, and a random effect to control for repeated samples and genetic relatedness. False discovery rate (FDR) was calculated following the q-value method of (Storey and Tibshirani, 2003). For the polyI:C data set, the observed p-value distributions were compared to an empirical null p-value distribution derived from re-running the same models but permuting the variable of interest (either condition or *CBD103* genotype) across 100 permutations of the data). For the CDV data set, the sample size was too small to calculate FDR based on the permutation-derived empirical null. We therefore estimated FDR by comparison to a uniform null, using the R package *qvalue* (Storey *et al.*, 2017) with the pfdr argument set to true. Because the assumption of a uniform null is likely anti-conservative for this analysis, we used more stringent criteria for significance by setting the FDR threshold to 1% and requiring that the absolute gene expression fold change be greater than 2.

To assess possible pathway or gene set enrichment among differentially expressed genes, we performed Gene Ontology enrichment analysis using g:ProfileR with the Ensembl 93 database (Raudvere *et al.*, 2019). We required a minimum of 5 and maximum of 350 genes in each tested biological process, and a minimum of 5 genes overlapping the biological process category and each of our gene sets of interest. We applied the “Best per parent group” hierarchical filtering option and used a Benjamini-Hochberg false discovery rate significance threshold of 0.01.

## Results

### Expression of CBD103 across tissues and cell types

To identify tissues where CBD103 is most likely to be involved in immune defense, we first assessed *CBD103* expression across a set of nine gray wolf tissues. *CBD103* was most highly expressed in epithelial tissues, including tongue and esophagus, consistent with expression patterns described for humans and dogs (Figure 1B) (Harder *et al.*, 2001; Leonard *et al.*, 2012). Based on published RNA-Seq data from whole blood (Charruau *et al.*, 2016) and newly generated RNA-Seq data for the two major skin cell types, keratinocytes and fibroblasts, *CBD103* has a unique expression pattern relative to other wolf beta defensins. Specifically, while all beta defensins we tested are very lowly expressed in fibroblasts and lowly expressed in whole blood, *CBD103* is uniquely highly expressed in keratinocytes (Figure 1C). These results led us to focus on challenge experiments in keratinocytes for the remainder of our study.

### Establishment of a panel of primary wolf keratinocytes

To assess the effects of *CBD103* genotype on wolf keratinocytes, we established a panel of primary wolf keratinocyte cultures. We first optimized conditions for growth of wolf keratinocytes (see Supplementary Materials and Methods). In brief, relative to three alternative conditions (FAD only, M199/M106 medium, or M199/M106 + RI), keratinocyte survival and proliferation rates were greatest when skin biopsies were plated in FAD medium and switched to FAD + RI two days post-plating, and supplemented with 3T3 feeder cells during serial passaging (Supplementary Figures S1-S3). One culture (from individual 15071) was used to estimate keratinocyte lifespan under these conditions and produced 28.2 cell population doublings over the course of 80 days (Supplementary Figure S5). We therefore used this protocol to culture keratinocytes from all subsequent samples (N = 24). The 24 keratinocyte cultures represented 14 and 10 wolves with the *K^yy^* and *K^yB^* genotypes, respectively (Supplementary Table S1).

### Establishment of a CBD103-edited wolf keratinocyte line

To isolate possible effects of the *CBD103 K^B^* allele on the keratinocyte response to virus, we immortalized a single *K^yy^* wildtype keratinocyte culture from individual 15071 (see Materials and Methods). We then used CRISPR/Cas9 editing to generate a line that was identical to the wildtype line, except for deletion of the three base-pair codon that distinguishes *K^y^* from *K^B^*, as well as a silent site substitution (G to A) incorporated as part of the gene editing process, and a single nucleotide intron substitution (C to G) that differentiates gray wolves from domestic dogs (because the donor oligonucleotide was designed based on the dog genome). Whole genome sequencing confirmed this sequence in and around the *CBD103* gene, as well as 323 potential off-target changes genome-wide (see Materials and Methods). Unexpectedly, however, RNA-Seq analyses revealed that *CBD103* expression was fully abolished in the edited *K^yB^* line relative to the original immortalized line (Supplementary Figure S6). Thus, we consider the *K^yB^* CRISPR/Cas9 edited line to function as a *CBD103* gene knockout line.

### Response of wolf keratinocytes to polyI:C

Given the role of *CBD103* in innate immunity (Leonard *et al.*, 2012), including in viral defense, we evaluated whether the *CBD103 K^y^/K^B^* genotype affects the gene expression response to a double stranded RNA virus mimic, polyI:C. To do so, we used RNA-seq to measure genome-wide gene expression in our 24 primary keratinocyte cultures, in two samples for each culture: (i) cells cultured in medium only (null); and (ii) cells exposed to 1 ug/mL polyI:C. PolyI:C elicited a strong gene expression response in wolf keratinocytes, such that the first principal component of overall gene expression levels separated null samples from polyI:C challenged samples (Figure 2A; R^2^ = 0.596, Pearson correlation p = 3.272 × 10^−10^). A total of 3,505 genes were up-regulated and 4,143 genes were down-regulated upon polyI:C stimulation (1% FDR; Supplementary Table S2). Genes up-regulated in response to polyI:C were enriched for 73 diverse biological processes, with the three most significantly enriched processes being “response to other organism”, “MAPK cascade”, and “inflammatory response” (all FDR < 1%), revealing the ability of primary wolf keratinocytes to mount an innate immune response to viral stimulation (Supplementary Table S3).

**Figure 2.**
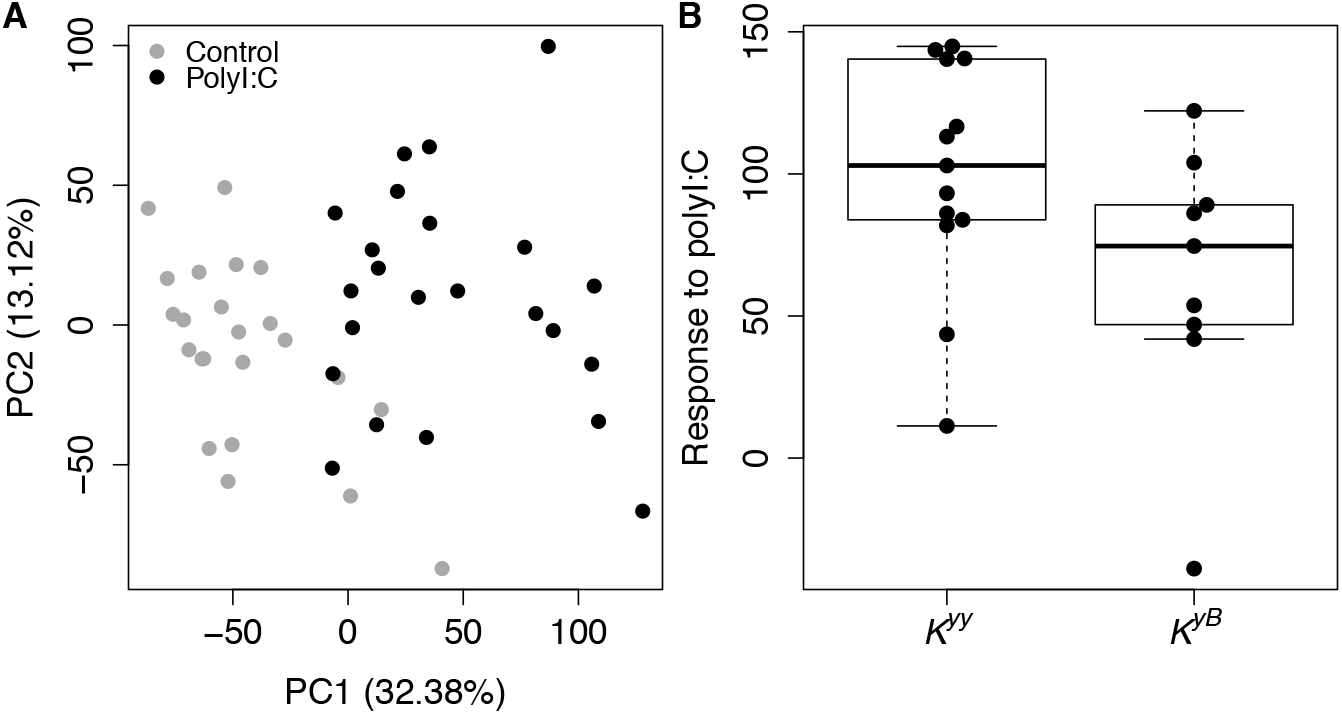
PolyI:C elicits a strong immune response in primary wolf keratinocytes that is not predicted by *CBD103 K* locus genotype. **(A)** Each dot represents a primary culture of wolf keratinocytes stimulated with polyI:C (black) or vehicle control (gray) for 24 hours. Values in parentheses in the x- and y-axis labels represent the percent of overall gene expression variation explained by principal components 1 and 2, respectively. PolyI:C treatment is strongly associated with PC1 of the overall gene expression data (R^2^ = 0.596, Pearson correlation p = 3.272 × 10^−10^). **(B)** The global gene expression response to polyI:C does not differ between *K^yy^* and *K^yB^* wolf keratinocytes (R^2^ = 0.154, Pearson correlation p = 0.070). Each dot represents the difference in PC1 between a matched pair of control and polyI:C stimulated samples. Each box represents the interquartile range, with a horizontal line depicting the median value. Whiskers indicate the most extreme values within 1.5x of the interquartile range.

In contrast, *CBD103* genotype was not associated with the overall response to polyI:C (as represented by the distance separating null and polyI:C samples for the same individual, on PC1 of the full gene expression data set: R^2^ = 0.154, Pearson correlation p = 0.070; Figure 2B). *CBD103* also did not predict gene expression for individual genes in either the null condition or in the polyI:C condition (all FDR > 10%; Supplementary Table S4). Finally, *CBD103* genotype did not predict individual gene responses to poly I:C (modeled as an interaction effect between *CBD103* genotype and polyI:C condition; 0 genes FDR < 1%, 7 genes FDR < 10%; Supplementary Table S5). This observation remained consistent even among genes that were most responsive to polyI:C stimulation (up-regulated by polyI:C at FDR < 0.1%; 0 genes FDR < 1%, 2 genes FDR < 10%; Supplementary Table S6). Thus, wolf keratinocytes carrying the *K^yB^* genotype appear to effectively be indistinguishable from keratinocytes with the *K^yy^* genotype in both null and polyI:C-stimulated conditions.

### Response of wolf keratinocytes to live canine distemper virus

The human homolog of CBD103 has been shown to have direct antiviral activity (Wilson *et al.*, 2013), and *CBD103* genotype is known to impact the binding affinity of the CBD103 peptide to other proteins (Candille *et al.*, 2007). We therefore also tested whether *CBD103* genotype affects the keratinocyte response to infection by live virus, using a GFP expressing strain of live canine distemper virus (CDV) (Von Messling *et al.*, 2004), an important pathogen in North American gray wolves (Almberg *et al.*, 2009). We observed that both primary keratinocytes and the immortalized *K^yy^* and *CBD103* gene knockout line are susceptible to infection, as evident by GFP expression within infected cells; increased rates of irregular cell shape, cell death and detachment; and up-regulation of key immune defense cytokines post-infection (Figure 3 and Supplementary Figures S7-S8).

**Figure 3.**
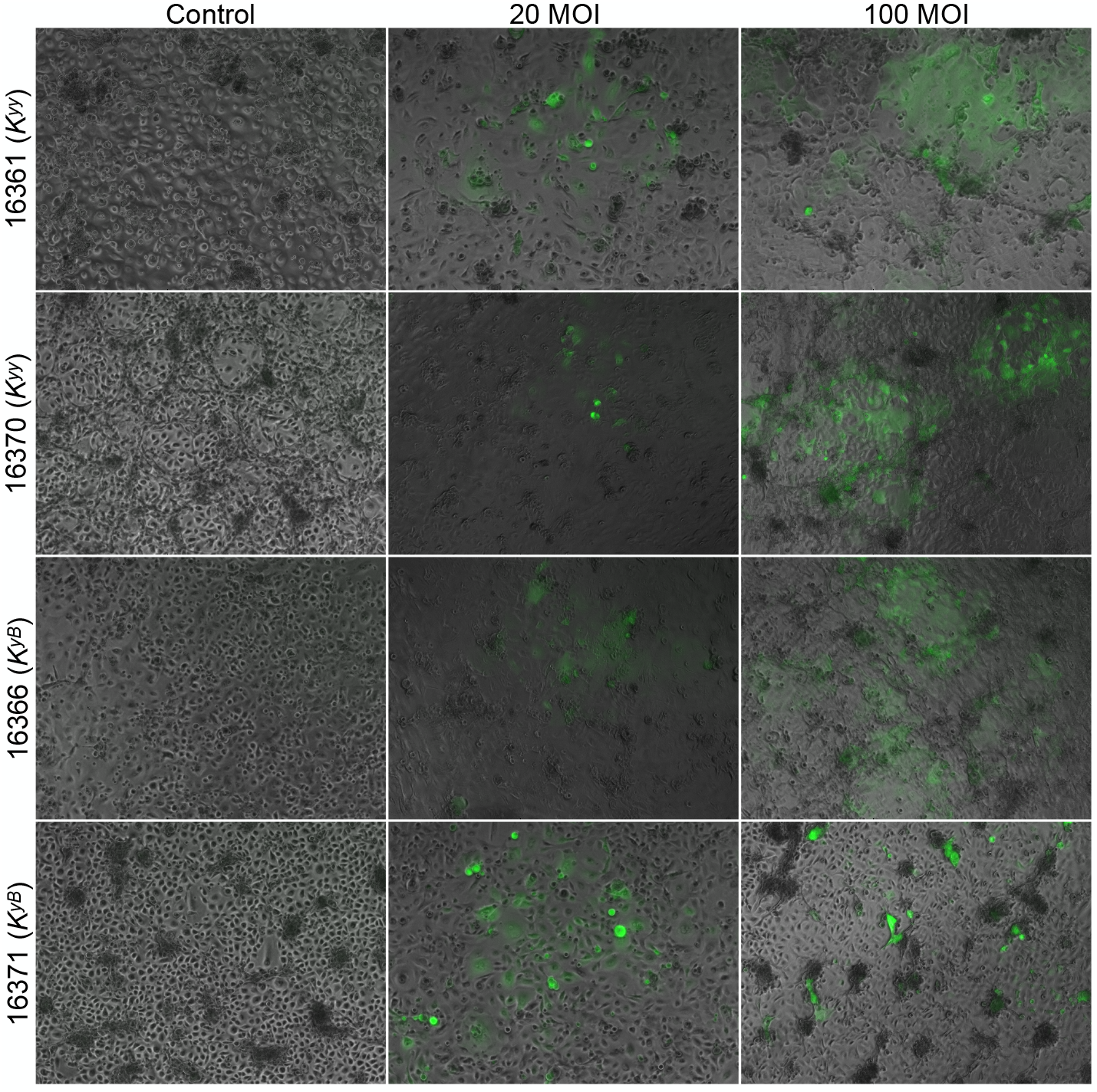
Primary wolf keratinocytes are vulnerable to canine distemper virus infection. *K^yy^* and *K^yB^* primary wolf keratinocytes infected with canine distemper virus for five days. Keratinocytes were infected at an MOI of 20 and 100 TCID_50_/cell. Fluorescence images were captured to visualize CDV (expressing GFP) and overlaid on phase contrast images. Animal IDs and corresponding *CBD103* genotypes are indicated on the y-axis.

We challenged a subset of primary keratinocyte cultures (N = 6; three *K^yy^* and three *K^yB^* primary cultures) with live CDV (MOI of 100 TCID_50_/cell) for five days. Live CDV elicited a moderate gene expression response in wolf keratinocytes, such that treatment condition (null versus CDV-challenged) was correlated with PC2 of overall gene expression (R^2^ = 0.397, Pearson correlation p = 0.028). A total of 348 genes were differentially expressed in response to infection by CDV (FDR < 1% and absolute fold change ≥ 2; Supplementary Table S7). The 156 genes up-regulated in response to CDV were enriched for “response to external biotic stimulus” (which included antiviral- and interferon-signaling genes *CXCL10*, *DDX60*, *DHX58*, *ISG15*, *ISG20*, *OAS1*, and *OAS3*) and “negative regulation of viral process” (FDR < 1%; Supplementary Table S8), suggesting that primary wolf keratinocytes are able to mount an innate immune response to infection by live CDV. The multi-day response to CDV, a live single-stranded RNA virus, provides distinct information from the 24 hour response to polyI:C, a double-stranded RNA virus mimic. Specifically, wolf genes that were responsive to CDV do not significantly overlap with those that were responsive to polyI:C, probably in part because the time points we used for measurement also differed (Fisher’s exact test OR (95% CI) = 1.007 (0.758, 1.351), p = 1).

*CBD103* genotype did not predict gene expression levels after CDV treatment (modeled as *CBD103* genotype nested in CDV; all FDR > 1%; Supplementary Table S9) or in response to CDV (modeled as an interaction term between *CBD103* genotype and CDV; all FDR > 1%; Supplementary Table S10), although our small sample size limits the power to detect such effects. Nevertheless, these results are congruent with those for polyI:C in suggesting that keratinocytes carrying the *K^yB^* genotype do not differ from *K^yy^* cells in the gene expression response to virus.

## Discussion

Previous research indicates that the *CBD103 K^y^*/*K^B^* polymorphism in North American gray wolves not only affects fur color, but has pleiotropic effects on other fitness-related traits (Coulson *et al.*, 2011; Stahler *et al.*, 2013; Cassidy *et al.*, 2017; Schweizer *et al.*, 2018). Most notably, *K^BB^* and *K^yB^* wolves have indistinguishable black fur color, but *K^BB^* wolves have much lower fitness than *K^yB^* wolves (Coulson *et al.*, 2011). Given that *CBD103* is a member of a gene family that is associated with innate immune function, we hypothesized that the *K^B^* allele might also alter immune defense, and specifically, the response to virus, in a manner that contributes to its intriguing evolutionary dynamics. Indeed, there is some evidence that immune function can be affected by pigmentation-related pathways. For example, the MC1R agonist alpha-melanocyte-stimulating hormone (α-MSH) also has microbicidal activity (Singh and Mukhopadhyay, 2014), and experimental manipulation of the amino acid sequence of α-MSH affects this activity (Grieco *et al.*, 2003). However, pleiotropic effects on immune function have yet to be explored for pigmentation variants that are naturally occurring.

Our data provide no strong support for the hypothesis that *CBD103* genotype alters the immune response to virus. *CBD103* genotype did not predict differences in the gene expression response to either polyI:C or to canine distemper virus, nor was it clearly associated with variation in baseline gene expression levels. Additionally, expression levels of *CBD103* itself did not differ depending on the *K* locus genotype. We note that, because our small sample size limits statistical power, particularly for the CDV data set, our results are limited to excluding the possibility that *CBD103* genotype effects are moderate to large. It therefore remains possible that *CBD103* has small effects on gene expression and/or the response to viral infection that would require larger sample sizes to detect.

Our findings suggest that, if the fitness consequences of *CBD103* genotype in wolves are indeed due to differences in immune function, they do not arise from simple differences in the immediate gene regulatory response to viruses in keratinocytes. However, *CBD103* genotype could influence immunity through mechanisms not evaluated here. First, the fitness effects of *CBD103* genotype could be mediated by viruses that we did not evaluate in this study, such as canine parvovirus or canine adenovirus-1 (Almberg *et al.*, 2009). Second, in humans, beta defensin 3 appears to have more pronounced antiviral activity in low salt concentrations similar to the environment of the oral cavity (Quiñones-Mateu *et al.*, 2003). Such concentrations would be lethal to the keratinocytes we studied here (derived from skin from ear punches), but could potentially be tested using epithelial cells established from the oral cavity or by incubating the virus with beta defensin 3 prior to cell infection (as in Quiñones-Mateu *et al*., 2003). Finally, given evidence that CBD103 can directly interact with pathogens (García *et al.*, 2001; Harder *et al.*, 2001; Wilson *et al.*, 2013), the codon deletion in the *K^B^* allele could alter the strength of direct protein-pathogen interactions. In the case of viruses like CDV, such a mechanism could affect viral entry rates, which could be tested by measuring intracellular viral titers over time. Cellular interactions with other types of pathogens or parasites known to affect North American gray wolf populations may also be relevant (e.g., wolves are vulnerable to *Sarcoptes scabiei-* induced sarcoptic mange: (Almberg *et al.*, 2009)). Importantly, the methods we applied here provide a basis for testing several of these possibilities *in vitro*.

Although our results do not resolve the mechanisms explaining the fitness costs of the *K^B^* allele in wolves, they do exemplify new approaches for studying genotype-phenotype relationships in non-model organisms. Specifically, we show that methods that were originally developed to culture primary human keratinocytes can also be applied to samples collected from a wild mammal population (Rheinwald and Green, 1975; Wu *et al.*, 1982). This approach makes it feasible to generate population samples of keratinocyte lines for experimental studies that cannot be conducted *in vivo*. The ability to produce a >2^25^-fold expansion of cells from small skin biopsies demonstrates the potential to cryopreserve many vials of early passage cells (on the order of >10^9^ total cells), permitting hundreds of experiments. Such methods could be useful for studying cellular responses to cutaneous pathogens, such as the white-nose syndrome epidemic that has devastated wild bat populations (Cryan *et al.*, 2010). With respect to pigmentation phenotypes, the cellular approaches we describe here could also be adapted for melanocyte culture (Costin *et al.*, 2004) or melanocyte-keratinocyte coculture (Lei *et al.*, 2002) to study the effects of genetic variation on the molecular basis of pigmentation phenotypes.

More broadly, cell culture methods provide a much needed approach for testing the explosion of hypotheses now being generated in evolutionary genomics. The reduced costs of high-throughput sequencing make genome-wide association studies, selection scans, transcriptomic studies, and comparative genomic analyses feasible in a broad spectrum of species. For example, analyses of selective sweeps in wolves have revealed many potential target genes, including those related to hypoxia and cold adaptation (Schweizer *et al.*, 2016a; Schweizer *et al.*, 2016b). However, genetic manipulations to functionally dissect these signatures are not ethical or feasible for most mammals, especially those that are declining or already rare or endangered. Yet many of these species can be sampled, non-lethally, for culturable cells and tissues such as skin, and live cells can even be collected from recently deceased individuals. Although fibroblast culture has long been used as a method of preserving genetic material (e.g., the “Frozen Zoo;” (Benirschke, 1984)), primary cell culture has not yet been widely applied to study inter-individual variation within or among populations. Such an approach is likely to become more attractive as immortalization, gene editing, and induced pluripotent stem cell (iPSC) generation becomes more feasible (e.g., (Ben-Nun *et al.*, 2011; Marchetto *et al.*, 2013; Romero *et al.*, 2015)). These approaches may be particularly valuable for isolating the causal effects of specific variants of interest, especially if those variants are rare (as in the case of *K^BB^*, estimated to be at 2% frequency in the Yellowstone gray wolf population) (Coulson *et al.*, 2011).

## Acknowledgments

We thank the wolf team at Yellowstone National Park, Jennifer Struthers, Jason Husseman, and their teams at the Idaho Department of Fish and Game for their hard work to collect samples for this project. We thank Jessica Cinkornpumin for guidance on lab work, and Steven Nguyen, Kashif Iqbal, Nadia Riabkova, and Joey Curtiss for their help with much of the molecular and cell work. We also thank Dr. Veronika von Messling, who kindly provided us the CDV 5804PEH-eGFP strain.

## Supplementary Material

Supplementary data are available at *Journal of Heredity* online.

## Data Availability

RNA sequencing data generated for this study are available in the NCBI Gene Expression Omnibus (GEO series accession GSE163163). RNA sequencing data from blood were previously published and available under GEO series accession GSE80440. DNA sequencing data of the immortalized cell lines are available in the NCBI Sequence Read Archive (BioProject accession number PRJNA688221).

