## Supplementary Materials and Methods, Figures for "Experimental assessment of K locus effects on the gray wolf response to virus"

**Table of Contents:**

**1. Supplementary Materials and Methods**

**2. Supplementary Figures**

Figure S1. Growth of wolf primary keratinocytes from skin biopsies across media conditions.
Figure S2. Growth of primary keratinocytes from a single wolf (16361) in different media.
Figure S3. Barplot of growth of wolf keratinocytes on different media.
Figure S4. Agarose gel electrophoresis to identify putative *CBD103* three base pair deletion clones.
Figure S5. Number of population doublings of primary keratinocytes established from a single gray wolf (individual 15071).
Figure S6. Expression of *CBD103* in the immortalized wolf keratinocyte lines.
Figure S7. Infection of immortalized wolf keratinocyte lines with live canine distemper virus (CDV).
Figure S8. Response of wolf keratinocytes to live CDV.

### Supplementary Materials and Methods

*Primary culture of wolf keratinocytes and fibroblasts*

To determine optimal growth conditions for wolf keratinocyte cell growth at initial plating, four skin biopsies were each split into two p60 Petri dishes. Two biopsies were fed FAD medium (1:1 DMEM:F12 base medium [ThermoFisher Scientific] + 5% iron-supplemented bovine calf serum [BCS; Hyclone, Cytiva Life Sciences] + 0.4 ug/ml hydrocortisone + 10 ng/ml epidermal growth factor [EGF] + 1% Penicillin-Streptomycin [ThermoFisher Scientific]) (Wu *et al.*, 1982), and two biopsies were fed M199/M106 medium (1:1 vol/vol M199:M106 medium [ThermoFisher Scientific] + 15% BCS + 10 ng/ml EGF + 0.4 ug/ml hydrocortisone + 1% Penicillin-Streptomycin) (Dickson *et al.*, 2000). After two days incubation, 10 uM ROCK inhibitor Y-27632 (hereafter referred to as RI; Cayman Chemical, Ann Arbor, MI) (Liu *et al.*, 2012) was added to one p60 of each of the paired cultures. We then quantified the number of skin pieces showing keratinocyte outgrowth after 4 days of incubation (Supplementary Figures S1-S3). All images were taken using an AxioCamMRm camera (Zeiss).

To determine the optimal medium for serially passaging the wolf keratinocytes, we compared the growth rates of keratinocytes cultured from individual 16361 and individual 16366 on three medium formulations: Keratinocyte serum-free medium (KSFM; ThermoFisher Scientific) + 25 µg/ml bovine pituitary extract (BPE; ThermoFisher Scientific) + 0.4 mM $CaCl_2$ + 0.2 ng/ml EGF + 1% Penicillin-Streptomycin (Dickson *et al.*, 2000), FAD medium, and

M199/M106 medium. We also tested the effect of including 10 uM RI in the medium and the effect of plating keratinocytes with 3T3 feeder cells, a mouse embryonic fibroblast line (Todaro and Green, 1963) that promotes the growth of human epidermal keratinocytes (Rheinwald and Green, 1975). Confluent cultures of 3T3 cells were treated with 3 μg/ml mitomycin-C for two hours and then were plated at 2 x $10^4$ cells/$cm^2$ together with the desired number of keratinocytes. This resulted in the comparison of 12 growth conditions. To quantify cell growth rate, we plated keratinocytes at 5,000 cells per well of a 6-well plate in each condition, and quantified the final number of cells after five days of incubation. The number of population doublings (PD) undergone by cells during each passage was calculated as $\log_2$(number cells at subculture/number cells plated). To determine the total replicative lifespan / expansion potential of the cells, cumulative PD during serial passage was plotted against total time in culture until senescence (Dickson *et al.*, 2000).

*Relatedness estimates*

To estimate a relatedness matrix, we performed genotyping on the RNA-Seq data using the Genome Analysis Toolkit (GATK) (McKenna *et al.*, 2010). We applied the GATK SplitNCigarReads function followed by indel realignment. We then performed base recalibration on genotypes with GQ ≥ 4 from initial genotyping of the full RNA-Seq dataset with the UnifiedGenotyper function. Final genotyping was performed with HaplotypeCaller. Variants that did not pass the following thresholds were removed: in GATK VariantFiltration: QUAL < 100, QD < 2, MQ < 35, FS > 30, HaplotypeScore > 13, MQRankSum < -12.5, ReadPosRankSum < -8; in vcftools (Danecek *et al.*, 2011): --max-missing-count 2 --minGQ 99 --min-alleles 2 --max-alleles 2 --min-meanDP 5 --remove-indels. Missing values were imputed with *beagle* (Browning and Browning, 2007). A relatedness matrix was generated from the filtered vcf file (25,459 variants) using the relatedness2 option in vcftools (Danecek *et al.*, 2011) which follows the method of (Manichaikul *et al.*, 2010).

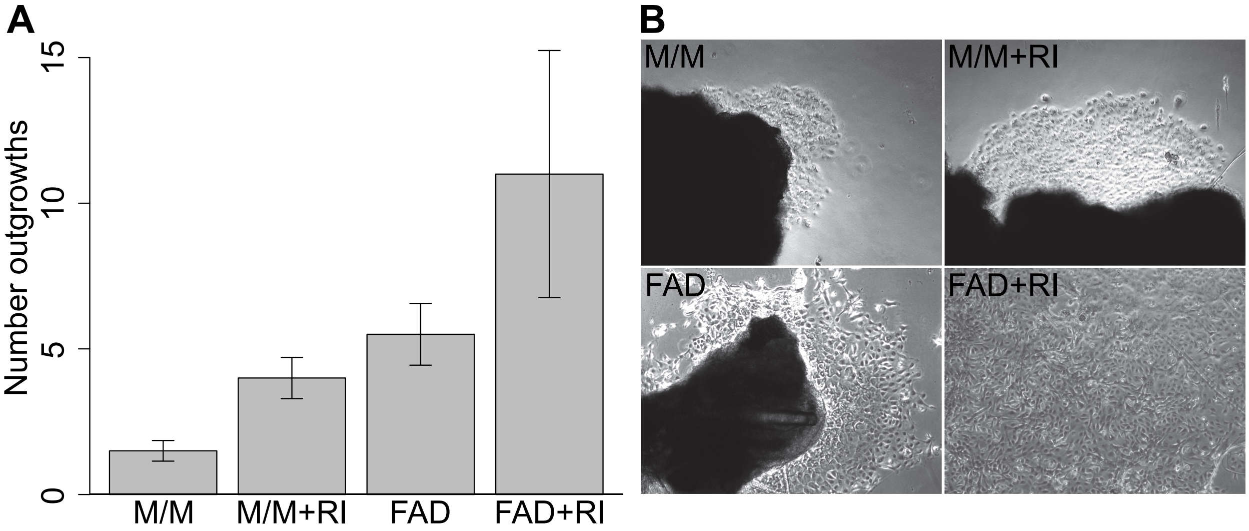


**Supplementary Figure S1**. **Growth of wolf primary keratinocytes from skin biopsies across media conditions**. Data and images collected six days after initial plating of minced 6 mm skin biopsies. **(A)** Number of outgrowths of keratinocytes from skin samples plated in M199/M106 (M/M) medium or FAD medium, with or without rock inhibitor (RI). Error bars represent the standard error of two skin biopsies (collected from different wolves) for each condition. **(B)** Keratinocytes in FAD medium exhibited healthier morphology (i.e., smaller, less flat, and fewer vacuoles) than keratinocytes in M199/M106 (M/M) medium, and keratinocytes in FAD medium with RI exhibited healthier morphology and more rapid proliferation than all three other treatments. Large dark areas in the first three panels are skin pieces.

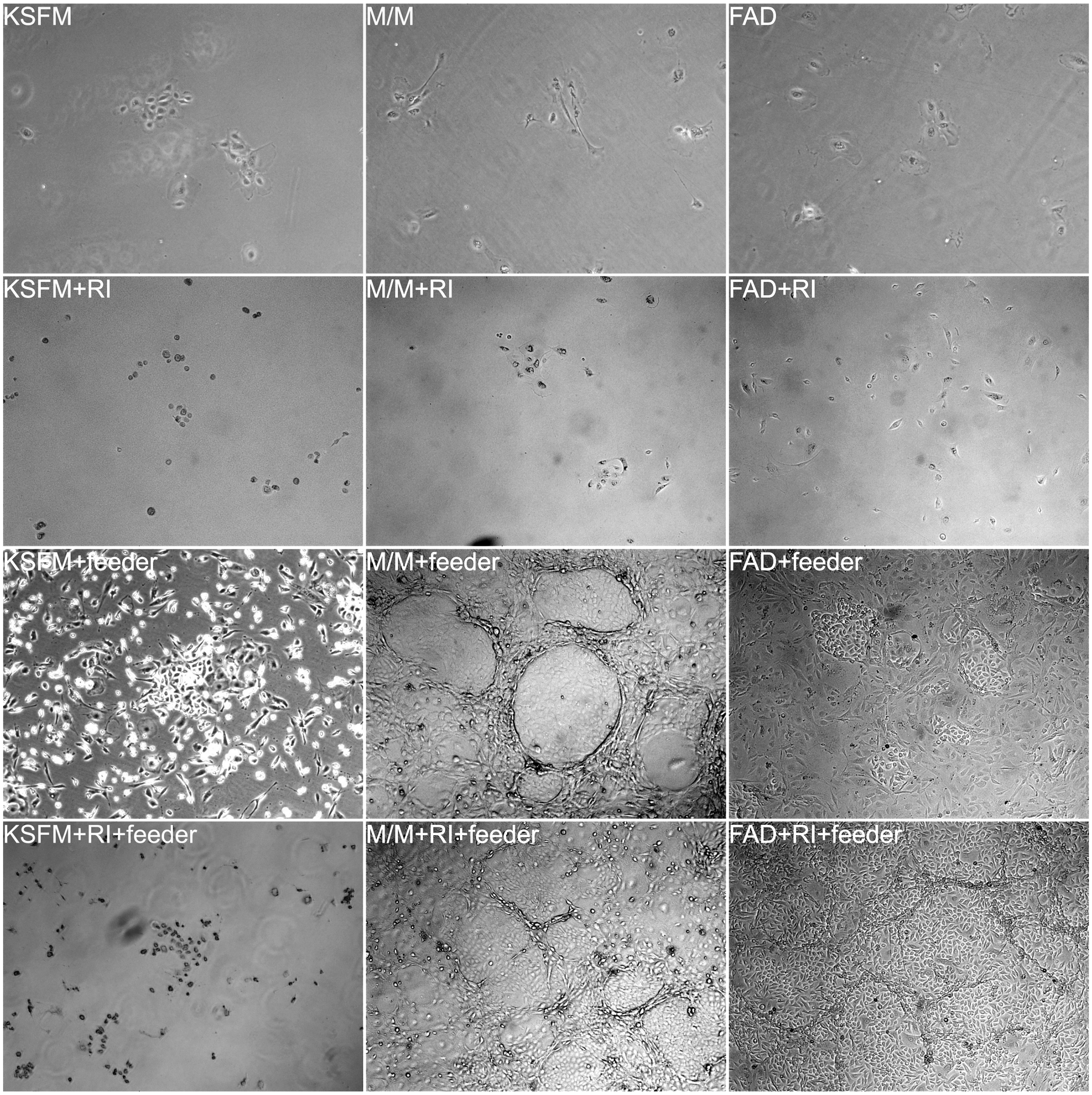


**Supplementary Figure S2**. **Growth of primary keratinocytes from a single individual (16361) in different media.** Keratinocytes were plated in parallel at 1 x $10^4$ cells per well in keratinocyte serum-free medium (KSFM), M199:M106 medium (M/M), or DMEM:F12 (FAD) medium with or without feeder cells or ROCK inhibitor (RI) (see Supplementary Materials and Methods for medium formulations). Images were taken 5 days after plating. Media with or without RI did not sustain keratinocyte proliferation, but the presence of 3T3 feeder cells improved keratinocyte health and proliferation in both M/M and FAD media (the feeder cells had a very low survival rate in KSFM). These results were further improved with addition of RI (see Supplementary Figure S3 for corresponding cell counts).

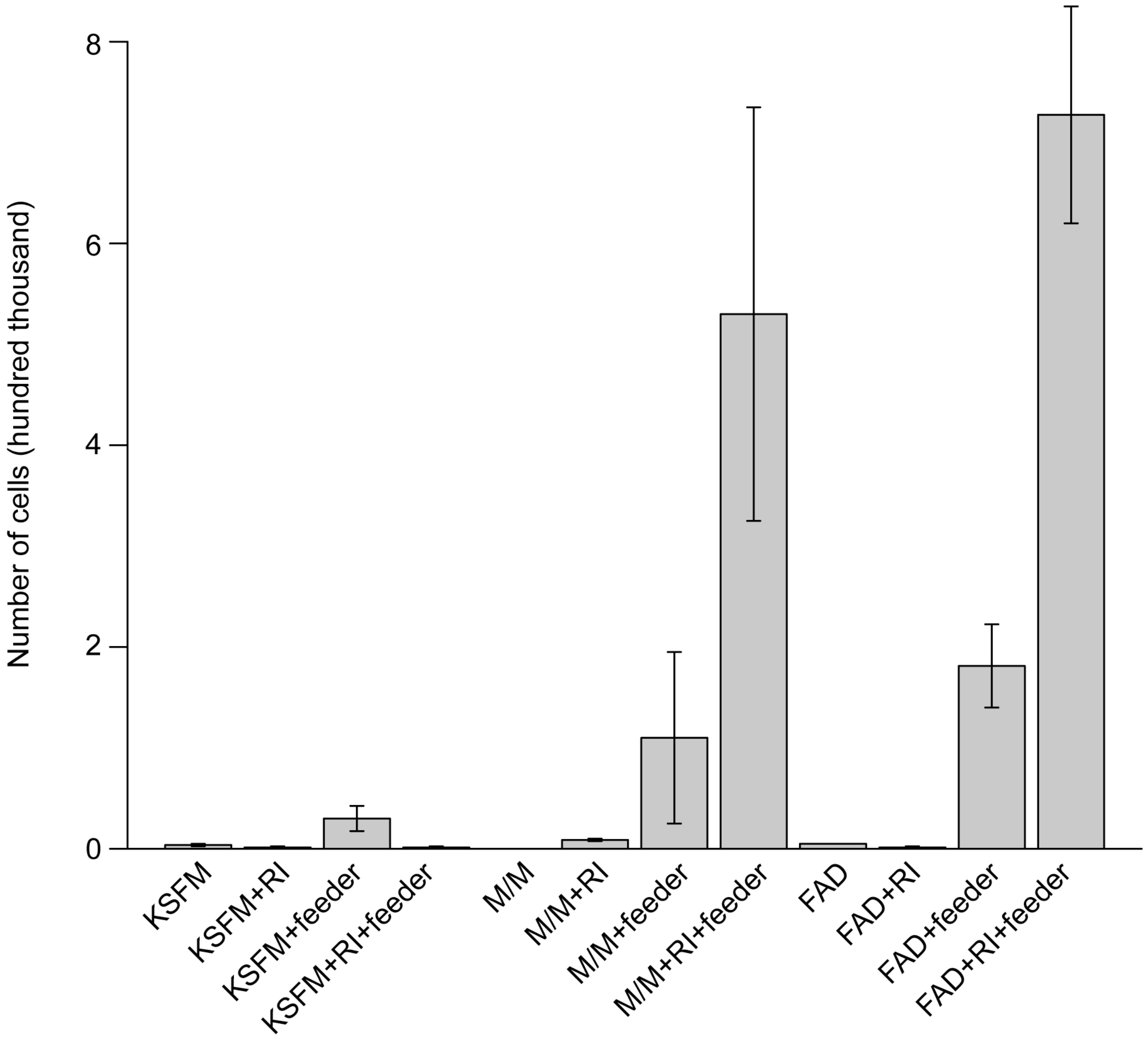


**Supplementary Figure S3**. **Barplot of growth of wolf keratinocytes on different media.** Keratinocytes were plated at 5,000 cells per well and cultured with different media for five days. Error bars represent standard errors of cells from two individual wolves (individuals 16361 and 16366). Abbreviations are the same as in Supplementary Figure S2.

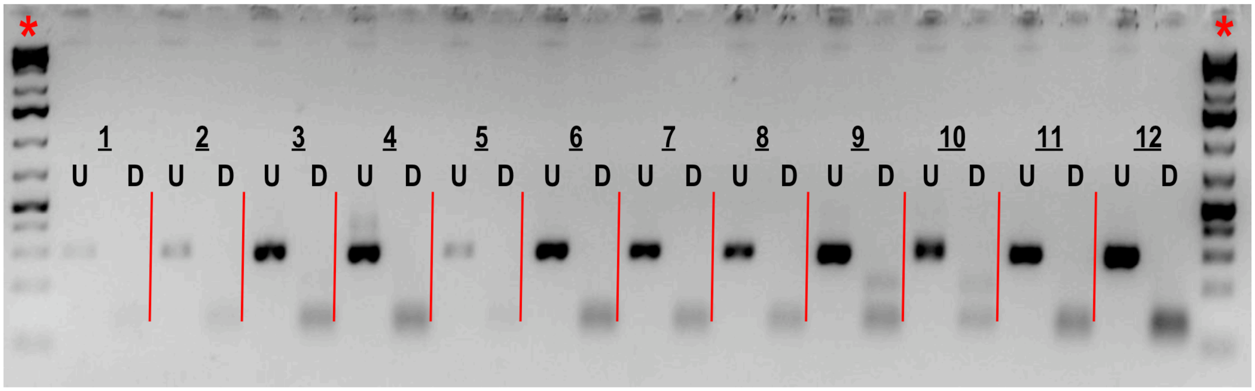


**Supplementary Figure S4. Agarose gel electrophoresis to identify putative *CBD103* three base pair deletion clones.** To identify clones that had putatively undergone successful CRISPR/Cas9 deletion of the three target *CBD103* base pairs (i.e., clones in which the $K^B$ allele was generated), single cell clones were expanded, and a 298 basepair DNA fragment surrounding the targeted three base pair deletion was PCR amplified. The PCR product was then digested with TspEI to distinguish between the $K^y$ and $K^B$ alleles. PCR product of $K^y$ yields three fragments of lengths 112 base pairs, 103 base pairs, and 83 base pairs, whereas TspEI digestion of the PCR product of $K^B$ yields two fragments, of lengths 112 base pairs and 183 base pairs. For each of the 12 clones depicted, band sizes of undigested DNA PCR product (U) and DNA product digested with TspEI (D) are shown side by side. Clone P2H9 (shown in lane 9) was identified as a putative *CBD103* three basepair deletion clone, and was confirmed by Sanger sequencing and by whole genome sequencing on the Illumina NovaSeq SP platform. Red lines separate lanes, and asterisks indicate DNA ladder.

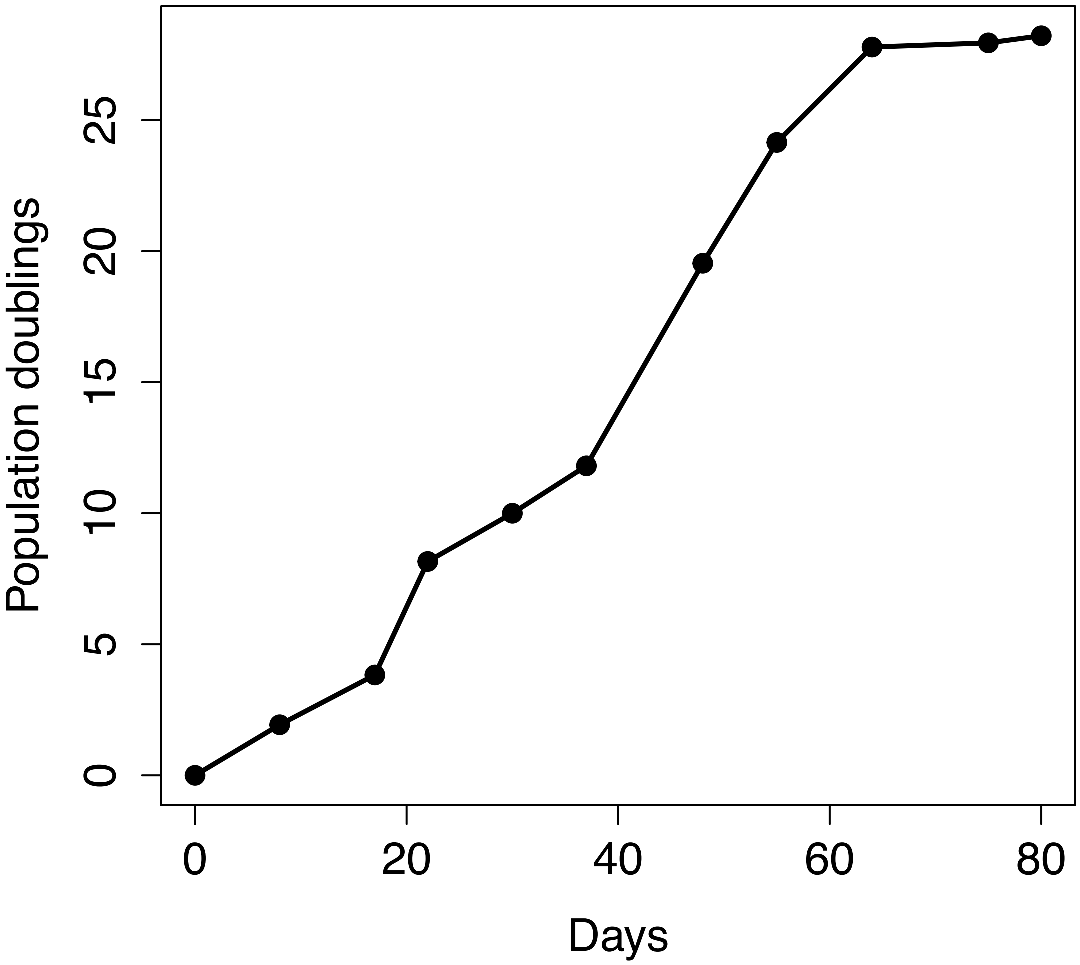


**Supplementary Figure S5**. **The total replicative lifespan of primary keratinocytes cultured from a single gray wolf (individual 15071).** Keratinocytes achieved a total of 28.2 population doublings after 80 days of culture in FAD medium with 3T3 feeder cells (with ROCK inhibitor). See Supplementary Materials and Methods for complete medium formulation.

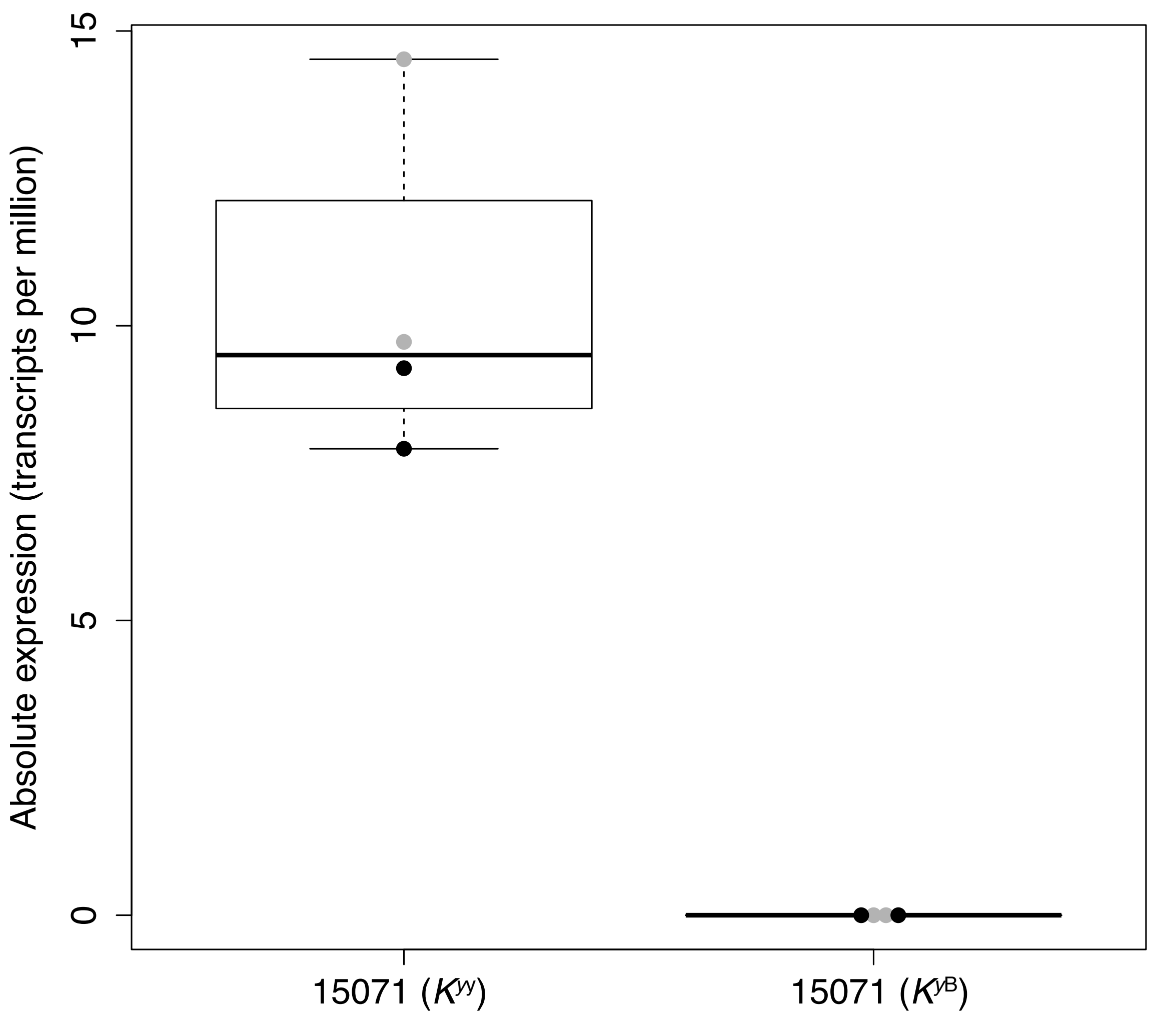


**Supplementary Figure S6**. **Expression of *CBD103* in the immortalized wolf keratinocyte lines.** Absolute expression of *CBD103* in the immortalized wildtype ($K^{yy}$) and CRISPR/Cas9 gene edited *CBD103* ($K^{yB}$) cell lines, measured with RNA-Seq. Each dot represents one well of cultured cells. Gray dots represent cells that were unstimulated, and black dots represent cells that were stimulated with polyI:C. Each box represents the interquartile range, with a horizontal line depicting the median value. Whiskers indicate the most extreme values within 1.5x of the interquartile range. *CBD103* expression was fully abolished in the $K^{yB}$ cell line (unpaired t-test, t = 7.197, df = 3, p = 0.006).

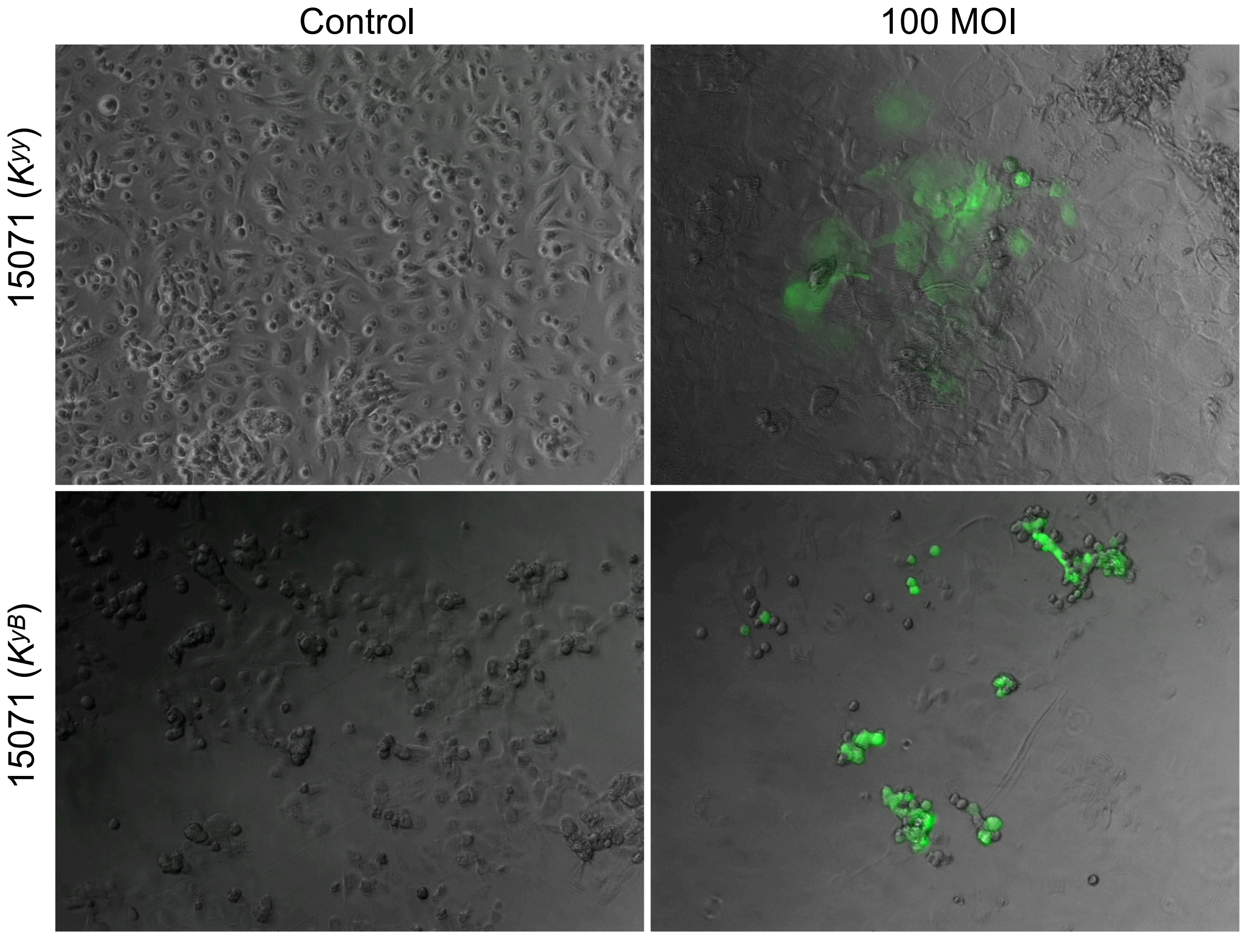


**Supplementary Figure S7**. **Infection of immortalized wolf keratinocyte lines with live canine distemper virus (CDV).** Immortalized wildtype ($K^{yy}$) and gene edited *CBD103* knockout ($K^{yB}$) keratinocytes infected with live CDV, five days post-infection. Keratinocytes were infected at an MOI of 100 $TCID_{50}$/cell. Fluorescence images were captured to visualize CDV (expressing GFP) and overlaid on phase contrast images.

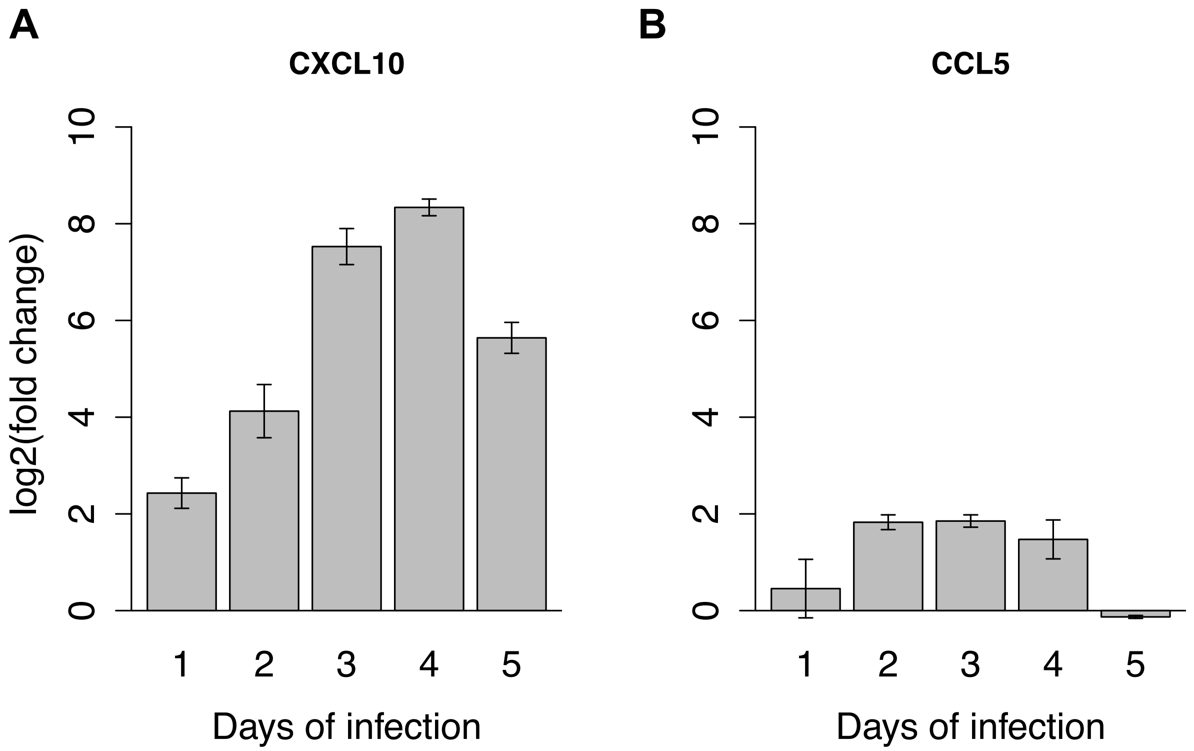


**Supplementary Figure S8**. **Response of wolf keratinocytes to live CDV.** Expression of *CXCL10* **(A)** and *CCL5* **(B)** of a single wildtype ($K^{yy}$) immortalized cell line in response to CDV at an MOI of 20 $TCID_{50}$/cell, 1 – 5 days post-infection relative to day 0. Error bars represent the standard errors of duplicate rt-qPCR measures. Note that when performing the CDV challenges for RNA-Seq, an MOI of 100 $TCID_{50}$/cell was used.
